# The Allosterically Modulated Free Fatty Acid Receptor 2 is Transactivated by an Increase in the Cytosolic Concentration of Calcium Ions

**DOI:** 10.1101/2022.10.15.512353

**Authors:** Simon Lind, Yanling Wu, Martina Sundqvist, Huamei Forsman, Claes Dahlgren

## Abstract

Allosterically modulated free fatty acid receptor 2 (FFA2R/GPR43) can be activated without the involvement of any orthosteric FFA2R agonist, by signals generated for example by P2Y_2_R, the G protein coupled receptor for ATP. An FFA2R specific positive allosteric modulator (PAM; Cmp58) was used to disclose the molecular mechanism by which signals generated by ATP/P2Y_2_R transactivates FFA2R. The P2Y_2_R induced signal that transactivates the allosterically modulated FFA2R was generated downstream of the Gα_q_ containing G protein that couple to P2Y_2_R. A receptor induced rise in the cytosolic concentration of ionized calcium ([Ca^2+^]_i_) was hypothesized to be the receptor transactivation signal. The Gα_q_ dependent transient rise in [Ca^2+^]_i_ induced by the ATP activated P2Y_2_Rs was not affected by Cmp58. The hypothesis gained, however, support from the finding that the modulator transferred FFA2R to a Ca^2+^ sensitive state. The rise in [Ca^2+^]_i_ induced by the Ca^2+^ specific ionophore ionomycin, activated the allosterically modulated FFA2R. The response induced by ionomycin was rapidly terminated and the FFA2Rs could then no longer be activated by the orthosteric FFA2R agonist propionate or be transactivated by the signal generated by the activated ATP receptor. The desensitized/non-responding state of FFA2R was, however, revoked by an earlier described cross-sensitizing/activating allosteric FFA2R modulator. The receptor transactivation of the allosterically modulated FFA2Rs, represent a unique regulatory receptor cross-talk mechanism by which the activity of a G protein coupled receptor is controlled by a signaling system operating from the cytosolic side of the plasma membrane.

**One-sentence summary:** A P2Y_2_ receptor signal generated downstream of a Gαq containing G protein transactivates the allosterically modulated FFA2 receptor

## Introduction

Neutrophils, innate immune defense cells and regulators of inflammatory reactions [1], are activated by various pathogen/danger associated molecular patterns (PAMPs/DAMPs) generated by microbes and/or injured host tissues. Such PAMPs/DAMPs are commonly recognized by neutrophil-expressed G protein coupled receptors [2, 3]. According to the classical recognition/signaling model, binding of a ligand to the orthosteric site in GPCRs transfers the information delivered from the external signaling molecule to the interior of the receptor expressing cell. This recognition and concomitant binding initiates receptor downstream signaling cascades cytosolically, and the signals generated regulate down-stream functional cell-responses [4]. The recognition/binding/signaling is no longer viewed as a simple on/off switch, with an activation state determined if the orthosteric binding pocket is empty or occupied by an agonist [5], and the results that have overturned this model include the finding that agonists can transduce functional selective cell responses. Such biased agonists preferentially trigger one signal-transduction pathway or functional response over another [5, 6]. The finding that also non-activating allosteric modulators affect receptor functions further increased the complexity of how GPCR signaling is regulated. Such modulators are recognized by receptor sites that are structurally and physically separated from the orthosteric binding pocket [7, 8] and binding of a modulator to such a site, changes the recognizing receptor allosterically and affects signaling induced by an orthosteric agonist either positively (amplify) or negatively (inhibit). This type of allosteric modulatory capability applies to many GPCRs, including the neutrophil free fatty acid receptor 2 (FFA2R; [9, 10]). Accordingly, the activity induced by short chain fatty acids, microbial/endogenous orthosteric FFA2R agonists, has been shown to be positively regulated by non-activating ligands that modulate function by binding to distinct allosteric receptor sites [11, 12].

Initially, the allosteric GPCR modulation concept was regarded to be receptor specific in that the modulation solely affected signaling induced by orthosteric agonists that bind to the same receptor as the modulator. This restriction is, however, not absolute [12, 13]. The allosterically modulated FFA2R may for example be transactivated by signals generated by another receptor, without the involvement of any orthosteric FFA2R agonist [11, 14, 15]. Receptor transactivation has been described for several receptor pairs and the mechanisms suggested include the generation of endogenous ligands for one receptor induced by signals generated by another receptor, and heterologous receptor phosphorylation by which an activated receptor modulates another receptor that concomitantly is turned to a signaling state [16–18]. The precise signaling mechanism at the molecular level, for how the transactivation of FFA2R is achieved, remains to be disclosed. When trying to dissect the molecular mechanisms underlying transactivation of FFA2R by signals generated down-stream of P2Y_2_R, the neutrophil receptor for ATP, we found that a rise in the intracellular concentration of free calcium ions ([Ca^2+^]_i_) was sufficient to activate allosterically modulated FFA2Rs. The data presented suggest that a rise in [Ca^2+^]_i_ is a key signal in transactivation of the allosterically modulated FFA2R, and this receptor activation is achieved without any orthosteric FFA2R agonist and controlled from the cytosolic side of the plasma membrane.

## Results

### The allosterically modulated FFA2R could be activated by two mechanisms that differ in sensitivity to Gα_q_ inhibition

In resting neutrophils, no activation of the superoxide (O_2_^−^) producing NADPH-oxidase was induced by the orthosteric FFA2R agonist propionate or the P2Y_2_R agonist ATP [15]. In the presence of the non-activating positive allosteric modulator (PAM) Cmp58, propionate but also ATP activated neutrophils to secrete O_2_^−^ (Fig 1A). In contrast to the propionate induced activity (Fig 1B and D), the response induced by ATP was inhibited both by AR-C118925, a P2Y_2_R specific antagonist, and by the Gα_q_ selective inhibitor YM-254890 (Fig 1A and C). In agreement with the involvement of FFA2R in the activation, the response induced by ATP, was also fully inhibited by CATPB (Fig 1), an FFA2R specific antagonist [19, 20]. The allosteric FFA2R modulator turned also nonactivating concentrations of PAF (recognized by the PAFR), fMLF (agonist for FPR1) and WKYMVM (agonist for FPR2) into activating agonists (Fig1A; inset).

**Fig 1.**
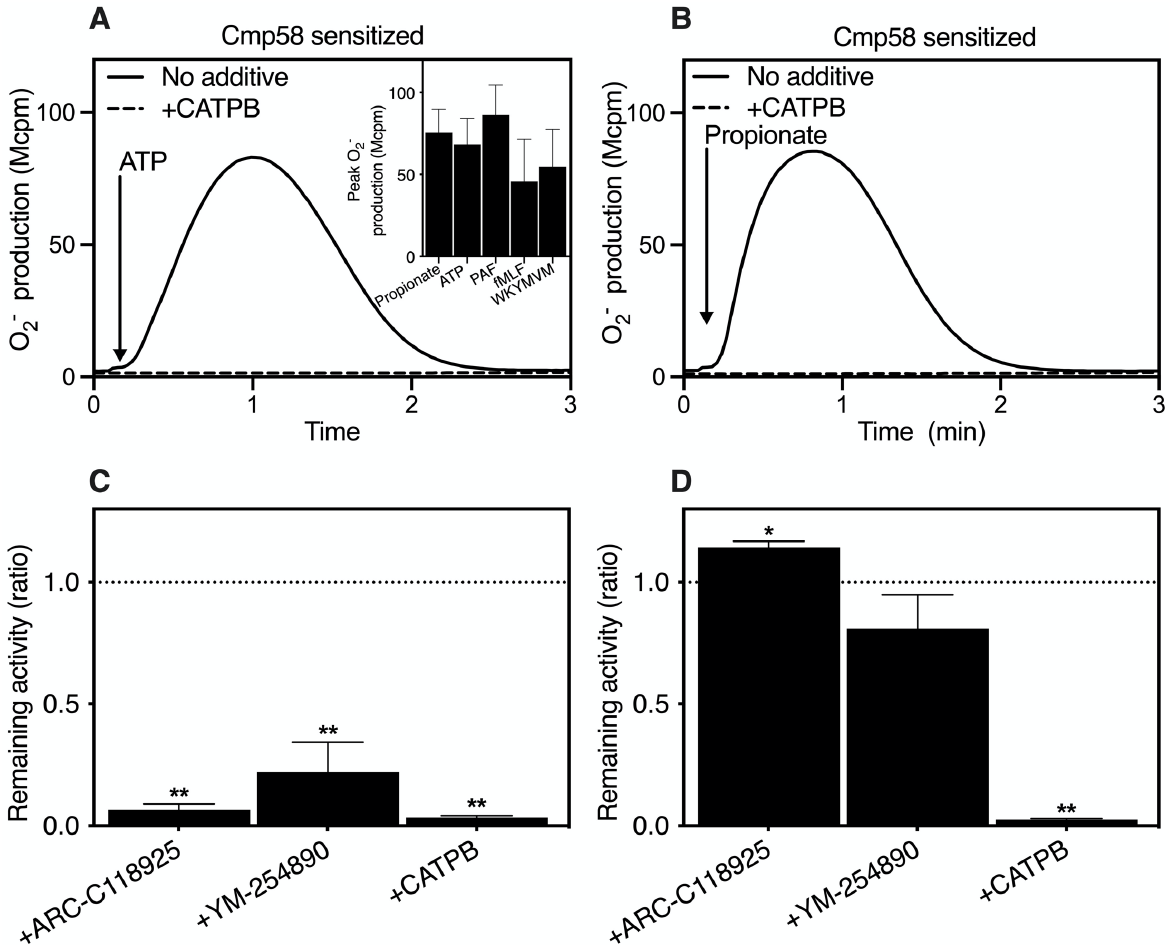
Neutrophil superoxide (O_2_^−^) production when activated by propionate and ATP, respectively -- effects of an allosteric FFA2R modulator, receptor antagonists and a Gα_q_ inhibitor. (**A**) Neutrophils preincubated with the allosteric FFA2R modulator Cmp58 (1 μM for 5 min) were activated by ATP (10 μM; time point for addition indicated by an arrow) in the absence (solid line) or presence (broken line) of the FFA2R specific antagonist (CATPB; 100 nM) and the production of O_2_^−^ was determined. One representative experiment out of >5 is shown. **Inset**: Neutrophils preincubated with Cmp58 (1 μM for 5 min) were activated by propionate (25μM), ATP (10 μM), PAF (1 nM), fMLF (0.5 nM) and WKYMVM (1 nM) and the production of O_2_^−^ was determined. The agonist concentrations were non-activating without Cmp58 and the responses were determined from the peak activities and expressed in Mcpm (mean + SD). (**B**) Neutrophils preincubated with the allosteric FFA2R modulator Cmp58 (1 μM for 5 min) were activated by propionate (25 μM; time point for addition indicated by an arrow) in the absence (solid line) or presence (broken line) of the FFA2R specific antagonist (CATPB; 100 nM) and the production of O_2_^−^ was determined. One representative experiment out of >5 is shown. (**C** and **D**) Neutrophils preincubated with the allosteric FFA2R modulator Cmp58 (1 μM for 5 min) were activated by ATP (10 μM; **C**) or propionate (25 μM; **D**) in the absence or presence of the P2Y_2_R specific antagonist ARC-C118925 (1 μM), the specific Gα_q_ inhibitor, YM-254890 (200 nM) and the FFA2R specific antagonist CATPB (100 nM) and the production of O_2_^−^ was determined. The responses were determined from the peak activities and presented as the ratio between the responses induced in the presence and absence of respective selective antagonist/inhibitor (mean +SD, n = 3). Statistical analyses were performed using a one-way ANOVA followed by a Dunnett’s multiple comparison test comparing the peak responses in the absence and presence of respective antagonist/inhibitor.

These data show that an activation of Cmp58-modulated FFA2Rs, can be achieved by two different mechanisms, and one of these (the ATP-P2Y_2_R signaling pathway) was achieved without involvement of any orthosteric FFA2R agonist and relied on signaling by a Gα_q_ containing G protein. Activation induced by the orthosteric FFA2R agonist propionate, was not affected by an inhibition of Gα_q_.

### The ATP/P2Y_2_R induced rise in intracellular Ca^2+^ was not affected by the allosteric FFA2R modulator Cmp58

Based on the fact that receptors shown to transactivate Cmp58-modulated FFA2Rs couple to Gα_q_ (i.e., the P2Y_2_R and PAFR) as well as to Gα_i_ (i.e., FPR1 and FPR2), some common receptor down-stream signal may be involved in the receptor cross-talk leading to a transactivation of FFA2R. It is well established that the Gα subunit of Gα_q_ containing G proteins as well as the Gßγ heterodimer subunit of a Gα_i_ coupled G protein can activate the PLC-PIP_2_-IP_3_-Ca^2+^ pathway [21] that results in a rise in the concentration of intracellular free Ca^2+^([Ca^2+^]_i_). This increase is initially obtained through a release of Ca^2+^ from intracellular stores and with some delay, by an opening of store-operated Ca^2+^ channels in the plasma membrane [22]. No rise in [Ca^2+^]_i_ was induced by the FFA2R PAM Cmp58 alone, whereas the modulator turned a non-activating concentration of propionate into an activation agonist. The propionate induced rise in [Ca^2+^]_i_ was fully inhibited by the FFA2R antagonist CATPB (Fig 2A). A transient rise in [Ca^2+^]_i_ was also induced by the P2Y_2_R agonist ATP alone but this response was not affected by Cmp58 (Fig 2A and B) and not inhibited by CATPB (Fig 2A).

**Fig. 2.**
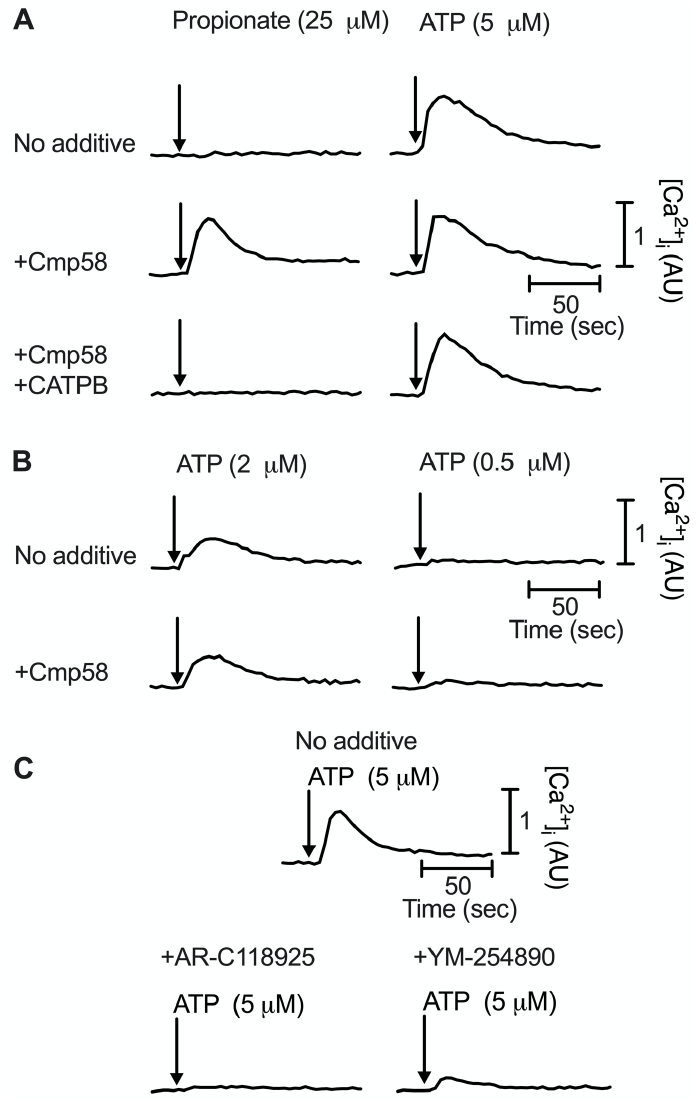
Neutrophil response, determined as a transient rise in the cytosolic concentration of free calcium ions ([Ca^2+^]), when activated by propionate and ATP, respectively -- effects of an allosteric FFA2R modulator, receptor antagonists and a Gα_q_ inhibitor. (**A**) Neutrophils preincubated without Cmp58 (No additive), Cmp58 alone (1 μM for 5 min; +Cmp58) or Cmp58 (1μM) combined with CATPB (100 nM; +Cmp58/+CATPB) were activated with propionate (25 μM; left panels) or ATP (5 μM; right panels). The change in [Ca^2+^]_i_. was followed and the time points for addition of propionate or ATP are indicated arrows. One representative experiment out of 3 independent experiment is shown. Abscissa, time of study (sec) Ordinate, the increase in [Ca^2+^]_i_ expressed as the ratio between Fura-2 fluorescence when excited at 340 nm and 380 nm, respectively. (**B**) Neutrophils preincubated without (No additive) or with Cmp58 (1 μM for 5 min; +Cmp58) were activated with ATP (2 μM; right panels or 0.5 μM; left panels). The change in [Ca^2+^]_i_. was followed and the time point for addition ATP is indicated by an arrow. One representative experiment out of 3 independent experiment is shown. Abscissa, time of study (sec) Ordinate, the increase in [Ca^2+^]_i_. expressed as the ratio between Fura-2 fluorescence when excited at 340 nm and 380 nm, respectively. (**C**) The inhibitory effects of the P2Y_2_R specific antagonist (+ARC-C118925; 1 μM) and the specific Gα_q_ inhibitor (+YM-254890; 200 nM) on the transient rise in [Ca^2+^]_i_ induced in neutrophils by ATP (5 μM; No additive). The time points for addition of ATP are indicated by arrows.

This ATP induced response was, however, fully inhibited both by the P2Y_2_R antagonist AR-C118925, and by the Gα_q_ selective inhibitor YM-254890 (Fig 2C).

These data show, that although the P2Y_2_R signals generated downstream of the Gα_q_ protein activate the allosterically modulated FFA2R, the P2Y_2_R generated transient rise in [Ca^2+^]_i_, was not affected by Cmp58. This suggests, that if a rise in the cytosolic concentration of Ca^2+^ is sufficient to activate the allosterically modulated FFA2R, the modulator affects the Ca^2+^ sensitivity of this receptor rather than signaling by the ATP/P2Y_2_R ligand/receptor complex.

### Ionomycin as well as thapsigargin activated FFA2Rprovided that the receptor was allosterically modulated by Cmp58

To study whether the allosteric modulator could indeed increase the sensitivity of FFA2R to a rise in the cytosolic concentration of Ca^2+^, we used the potent a selective calcium ionophore (ionomycin) as a tool compound. Ionomycin facilitates the entry of Ca^2+^ over biological membranes and increases the [Ca^2+^]_i_ by a mechanism that bypasses cell surface receptors. In agreement with earlier findings [23, 24], ionomycin when used as a neutrophil activating compound, increase the [Ca^2+^]_i_ and this increase was not affected by FFA2R ligands or accompanied by any pronounced secretion of O_2_^−^ (Fig 3) generated by the plasma membrane localized NADPH-oxidase. Cmp58 turned, however, not only propionate and ATP into potent oxidase activating agonists, but also ionomycin was turned into a potent neutrophil activator, determined as the release of O_2_^−^ (Fig 4A). This neutrophil response was totally inhibited by the FFA2R antagonist CATPB (Fig 4A and B) and dependent on the concentration of ionomycin used to induce a rise in [Ca^2+^]_i_ (Fig 4C). In contrast to the reciprocal relation between the allosteric FFA2R modulator Cmp58 and the orthosteric agonist propionate [15], the order by which the modulator and ionomycin was added to the neutrophils could not be reversed. That is, no activation of the oxidase is obtained when Cmp58 is added to ionomycin sensitized neutrophils (Fig 4D). The lack of reversibility was evident also when the FFA2R modulator was combined with ATP (Fig 4D).

**Fig 3.**
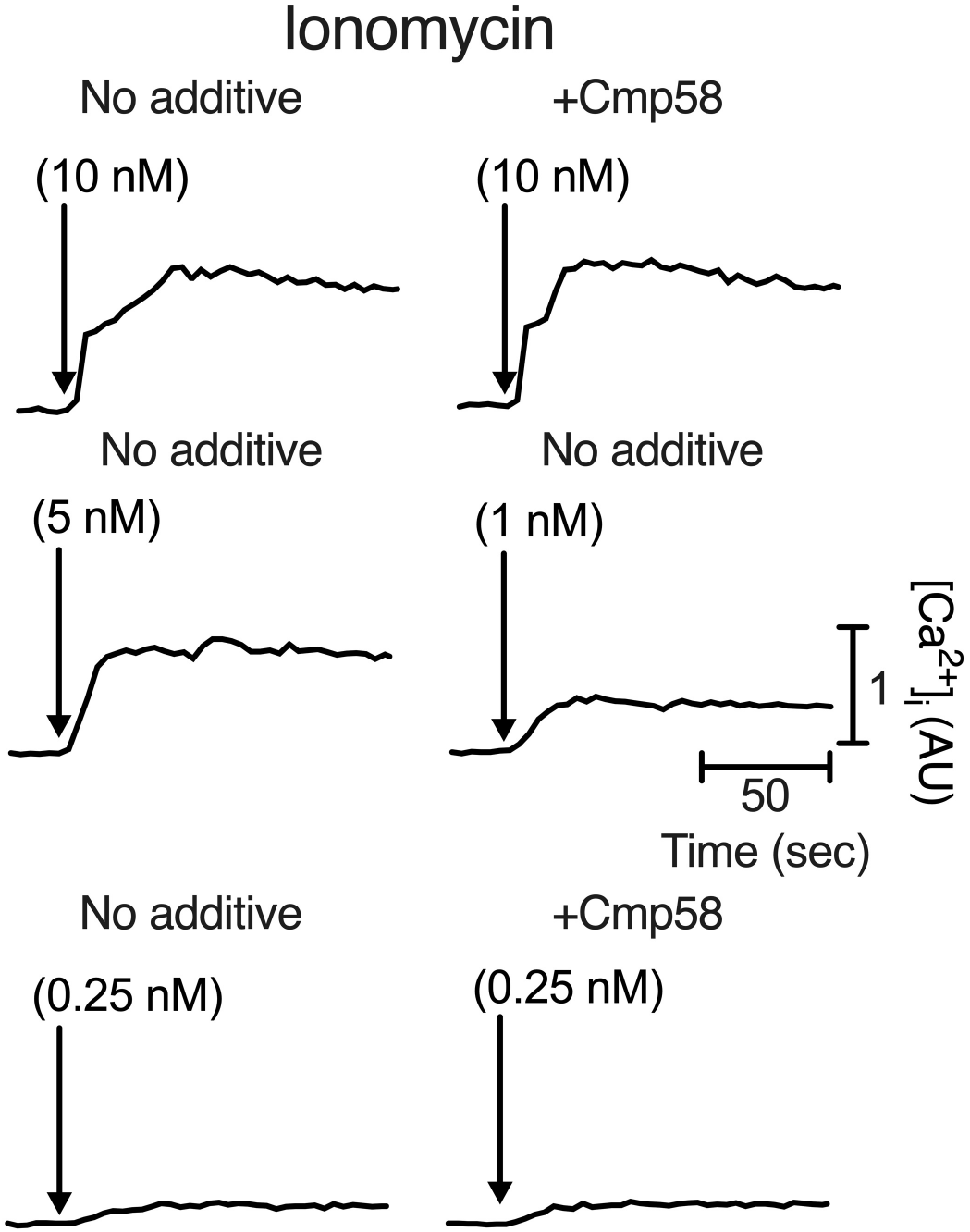
The ionomycin induced rise in the [Ca^2+^]_i_) is unaffected by the allosteric FFA2R modulator Cmp58. Neutrophils preincubated without (No additive) or with Cmp58 (1 μM for 5 min; +Cmp58) were activated by different concentrations of ionomycin (10, 5 and, 0.25 nM, respectively). The time points for addition of ionomycin are indicated by arrows. The change in [Ca^2+^]_i_ was followed and one representative experiment out of 3 independent experiment is shown. Horizontal bar; time of study (sec). Vertical bar; increase in [Ca^2+^]_i_ expressed as the ratio between Fura-2 fluorescence when excited at 340 nm and 380 nm, respectively.

**Fig 4.**
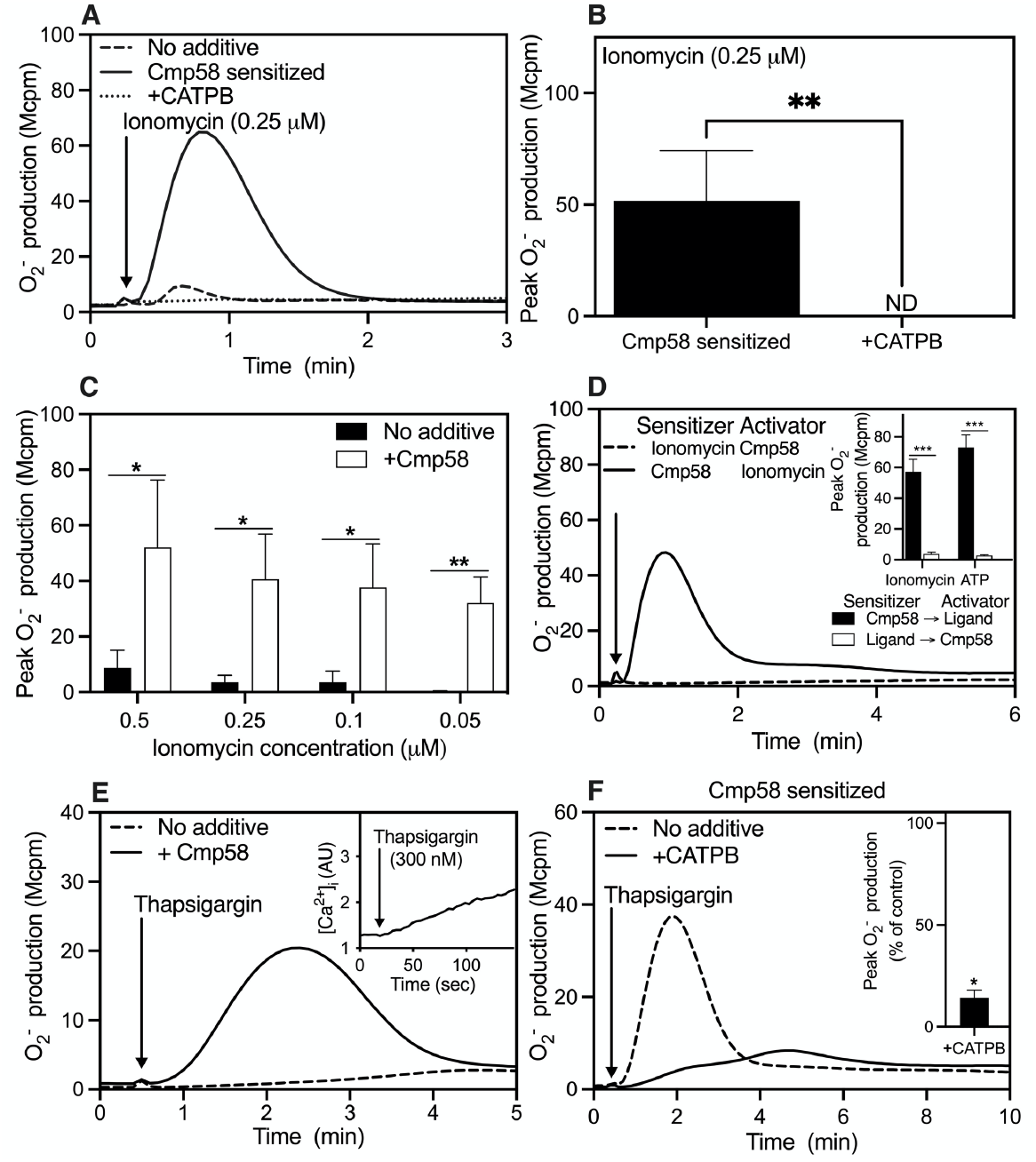
A rise in the [ca^2+^]_i_ potently activate neutrophils provided that FFA2R is allosterically modulated. (**A**) Neutrophils were incubated without Cmp58 (broken line) and with Cmp58 alone (1 μM for 5 min; solid line) or with Cmp58 (1μM) combined with CATPB (100 nM; dotted line) and the cells were then activated by ionomycin (0.25 μM; time point for addition indicated by an arrow). The production of (O_2_^−^) was followed and expressed in Mcpm. One representative experiment out of >5 is shown. (**B**) Peak O_2_^−^ production induced by ionomycin (0.25 μM) was determined in neutrophils sensitized with Cmp58 alone (1 μM, 5 min) or when combined with the FFA2R antagonist CATPB (100 nM). The responses were determined from the peak activities and expressed in Mcpm (mean+SD, n = 5). The statistical analysis was performed using paired Student’s t-test comparing the peak responses induced either by absence or presence of the antagonist CATPB. ND = not detectable (**C**) Production of O_2_^−^ in neutrophils activated with different concentrations of ionomycin in the absence (black bars) or presence of the FFA2R specific allosteric modulator Cmp58 (1 μM, 5 min; white bars,) respectively. The responses were determined from the peak activities and expressed in Mcpm (mean + SD, n = 3). The statistical analysis was performed using paired Student’s t-test comparing the peak responses induced either in absence or presence of Cmp58, respectively. (**D**) Neutrophils were incubated (sensitized) with ionomycin (0.25 μM, 5 min; broken line) or with Cmp58 (1 μM, 5 min; solid line). The ionomycin sensitized neutrophils were activated with Cmp58 (1 μM; time point for addition indicated by an arrow) whereas the Cmp58 sensitized neutrophils were activated by ionomycin (0.25 μM). The production of O_2_^−^ was measured and one representative experiment out of 3 is shown. **Inset**: Determination of the order by which ionomycin and Cmp58 was added to the neutrophils. The production of O_2_^−^ peak values was determined using the setup described above, as well as in the same setup with ATP (10 μM) replacing ionomycin. The responses were determined from the peak activities and expressed in Mcpm; mean +SD, n=3. (**E**) Neutrophils were incubated without Cmp58 (broken line), with Cmp58 alone (1 μM for 5 min; solid line) or with Cmp58 (1μM) combined with CATPB (100 nM; dotted line) and the cells were then activated by thapsigargin (300 nM; time point for addition indicated by an arrow). The production of (O_2_^−^) was followed and expressed in Mcpm. One representative experiment out of >5 is shown. **Inset**: The change in [Ca^2+^]_i_. induced in neutrophils by thapsigargin (300 nM) was followed. The time point for addition of thapsigargin is indicated by an arrow. One representative experiment out of 3 independent experiment is shown. Abscissa, time of study (sec) Ordinate, the increase in [Ca^2+^]_i_. is expressed as the ratio between Fura-2 fluorescence when excited at 340 nm and 380 nm, respectively. (**F**) Neutrophils incubated (sensitized) with Cmp58 alone (1 μM for 5 min; broken line) or with Cmp58 (1 μM) combined with the FFA2R specific antagonists, CATPB (100 nM; solid line). were activated by thapsigargin (300 nM dashed line; time point for addition indicated by an arrow) and the release of O_2_^−^ was measured continuously and expressed in Mcpm. **Inset**: Inhibitory effect of the FFA2R antagonist CATPB on the thapsigargin induced neutrophil O_2_^−^ production expressed in remaining activity (percent determined from the peak values of the responses; mean + SD, n=3). The statistical analysis was performed using paired Student’s t-test comparing the peak responses induced in the absence or presence of the antagonist CATPB, respectively, when activating with thapsigargin.

A selective inhibitor of a Ca^2+^-ATPase, an enzyme expressed in the Ca^2+^ storing endoplasmic reticulum, can also be used as a tool compound to increase the [Ca^2+^]_i_ without the direct involvement of any cell surface receptor. Thapsigargin is such an inhibitor that increases the [Ca^2+^]_i_ by an emptying of the intracellular stores [25]. Compared to ionomycin, a slower increase in [Ca^2+^]_i_ was induced by thapsigargin (Fig 4E; inset) and this increase alone was not linked to any activation of the NADPH-oxidase. More importantly, Cmp58 turned also thapsigargin into an activating compound determined as the release of O_2_^−^ (Fig 4E), and also this response was inhibited by the FFA2R antagonist CATPB (Fig 4F).

### Cmp58 could be replaced by the allosteric FFA2R modulator AZ1729 but not by the orthosteric FFA2R agonist propionate

In order for propionate to activate the neutrophil O_2_^−^ generating oxidase, FFA2R had to be allosterically modulated (see above). The non-activating FFA2R selective PAM AZ1729 has earlier been shown be recognized by an allosteric binding site physically/functionally separated from the binding site for Cmp58 [11, 26]. The presence of two different allosteric binding sites was evident by the fact that when combined, the non-activating allosteric FFA2R modulator Cmp58 turned the other modulator AZ1729 into a potent neutrophil activating ligand (Fig 5A). When AZ1729 was included as the allosteric FFA2R modulator in the receptor cross-talk/transactivation assay system, ATP was turned into an activating agonist (Fig 5B). The activation pattern in the presence of AZ1729, regarding the effects of receptor antagonists and Gα_q_ inhibition was identical to that for Cmp58 (see above).

**Fig 5.**
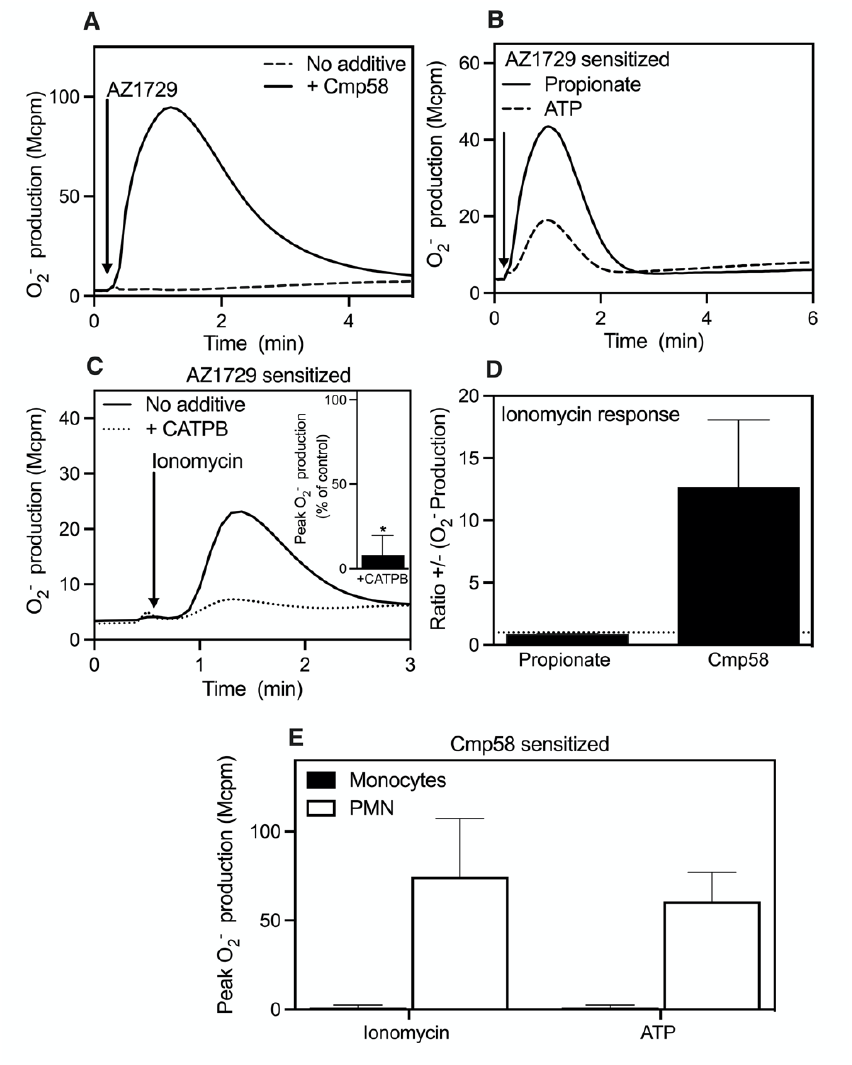
Cmp58 could be replaced by the interdependent allosteric FFA2R modulator AZ1729 but no activity is induced in monocytes. (**A**) Neutrophils incubated (sensitized) without (broken line) or with Cmp58 (100 nM; 5 min; solid line) were activated by the interdependent allosteric FFA2R modulator AZ1729 (1μM) and the production/release of O_2_^−^ was determined and expressed in Mcpm. One representative experiment out of >5 is shown, and the time point for addition of the agonists is marked by an arrow. (**B**) Neutrophils sensitized with AZ1729 (1 μM, 5 min) were activated by propionate (25 μM; solid line) or ATP (10 μM; broken line) and the production/release of O_2_^−^ was determined and expressed in Mcpm. One representative experiment out of >5 is shown, and the time point for addition of the agonists is marked by an arrow. (**C**) Neutrophils sensitized with AZ1729 alone (1 μM, 5 min; solid line) or combined with the FFA2R antagonist CATPB (100 nM, dotted line) were activated by ionomycin (0.25 μM; time point for addition indicated by an arrow) and the production/release of O_2_^−^ was determined and expressed in Mcpm. One representative experiment out of 3 is shown. **Inset.** Neutrophils preincubated with the allosteric FFA2R modulator AZ1729 (1 μM for 5 min) were activated by ionomycin (0.25 μM) in the absence or presence of the FFA2R specific antagonist CATPB (100 nM) and the production of O_2_^−^ was determined. The responses were determined from the peak activities and presented as the remaining activity (in percent) in the presence of the antagonist (mean + SD, n = 3). The statistical analysis was performed using paired Student’s t-test comparing the peak responses induced in the absence and presence of the antagonist. (**D**) Neutrophils incubated (sensitized) with propionate (25 μM; 5 min) or Cmp58 (1 μM; 5 min) were activated by ionomycin (0.25 μM) and the production/release of O_2_^−^ was determined and shown as fold increase determined from the peak activities in the absence and presence of the sensitizing compound (mean +SD, n=3). (**E**) Neutrophils (white bars) and monocytes (black bars) were incubated (sensitized) with Cmp58 (1 μM; 5 min) were activated by ionomycin (0.25 μM) or ATP (10 μM) and the production/release of O_2_^−^ was determined as the peak activities (Mcpm; (mean + SD, n=3).

AZ1729 turned also ionomycin to a neutrophil activating compound, and this response was totally inhibited by the FFA2R antagonist CATPB (Fig 5 C). No such activation was induced by ionomycin (or ATP) when the allosteric modulator was replaced by the orthosteric FFA2R agonist propionate (Fig 5D).

The same experimental receptor transactivation protocol as that used for neutrophils was transferred to monocytes, cells earlier shown not to express functional FFA2Rs [27, 28]. Monocytes do, however, respond to ATP with a transient rise in [Ca^2+^]_i_, thus showing that the cells express functional P2Y_2_Rs [15]. No NADPH oxidase activity was induced in monocytes by ATP or the calcium ionophore ionomycin, irrespectively if Cmp58 was present or not (Fig 5E). Taken together, these results show that in order for ATP and ionomycin to activate the NADPH oxidase system, functional FFA2Rs are required.

### No signaling role of granule produced reactive oxygen species

Despite the fact that very small amounts of superoxide anions were released by ionomycin-activated neutrophils (see above) the Ca^2+^ionophore has been shown to induce NADPH-oxidase dependent production of reduced/reactive oxygen species (ROS) intracellularly [23, 29]. Being highly reactive, such ROS have been suggested to possess signaling properties in GPCR transactivation of other receptors such as that recognizing epidermal growth factor [30]. Despite the fact that ionomycin induced a pronounced release of ROS in the presence of Cmp58 (see above), the intracellular oxidase activity induced by ionomycin was not affected by the allosteric FFA2R modulator or by the FFA2R antagonist CATPB (Fig 6A).To determine the signaling role (if any) of intracellularly produced ROS, in the allosterically FFA2R modulator dependent release of O_2_^−^ induced by ionomycin, we took advantage of the fact that the widely used pharmacological NADPH-oxidase inhibitor diphenyleneiodonium (DPI; [31]), when present in low micromolar concentrations, selectively inhibits intracellular (granule localized) ROS production [32]. In accordance with this, we show that the ionomycin induced intracellular ROS production was totally inhibited by DPI, whereas this inhibitor had no effect on the allosteric FFA2R modulator dependent release of ROS induced by ionomycin (Fig 6B). It should also be noticed that the kinetic difference between the intracellular and extracellular ROS production, respectively, support the conclusion that the granule produced ROS have no signaling role in the allosteric FFA2R modulator dependent release of ROS (Fig 6C). At the time point for the maximum ROS release (less than one min), the intracellular ROS production induced by ionomycin had just been initiated.

**Fig 6.**
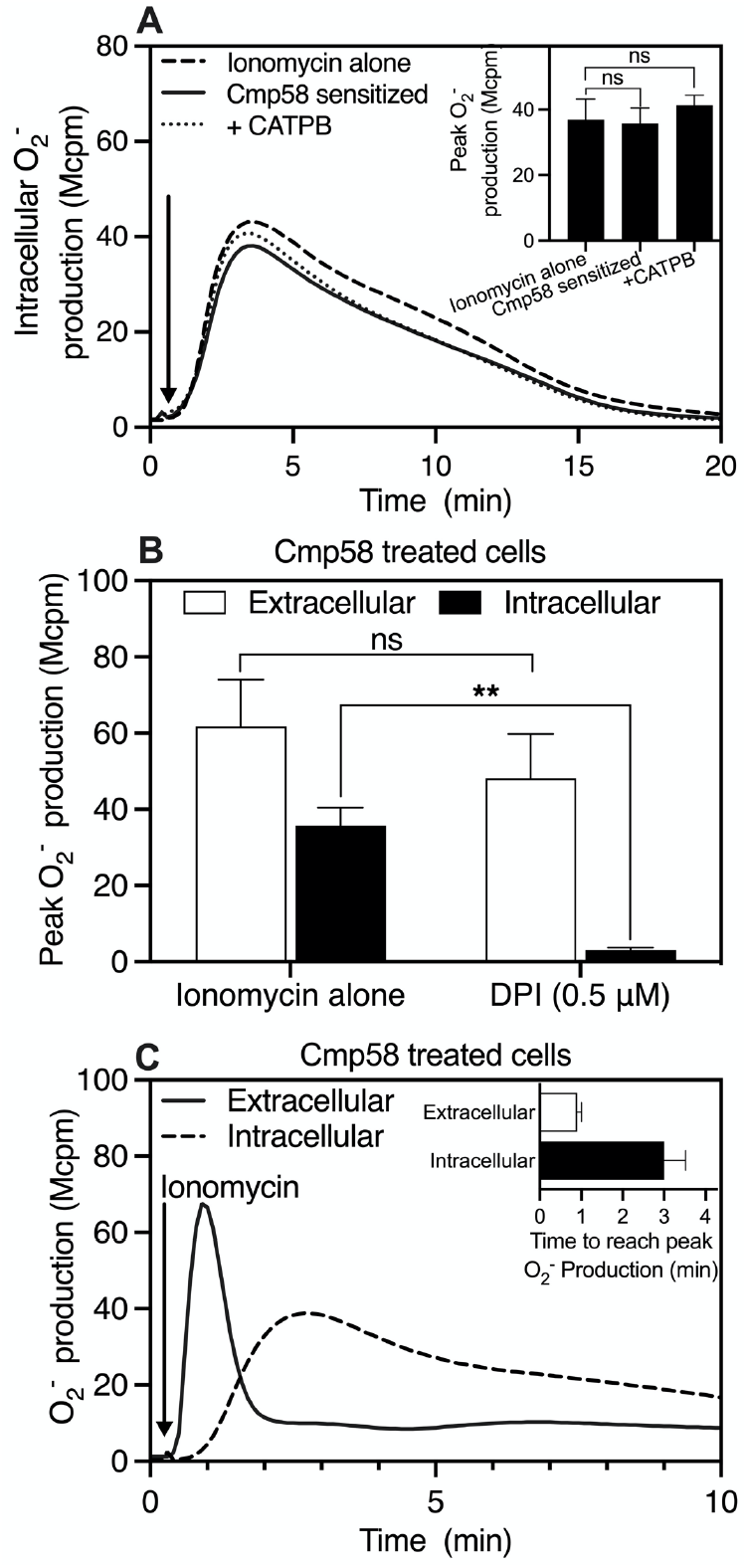
The FFA2R independent intracellular ROS production is inhibited by the NADPH oxidase inhibitor DPI – the FFA2R dependent release of ROS is not affected by DPI. (**A**) Neutrophils were incubated without Cmp58 (broken line), with Cmp58 alone (1 μM for 5 min; solid line) or with Cmp58 combined with CATPB (100 nM; dotted line) and the cells were then activated by ionomycin (0.25 μM; the time point for addition of the ionophore is indicated by an arrow). The production of intracellular (O_2_^−^) was followed and expressed in Mcpm. One representative experiment out of 3 is shown. **Inset**: Peak intracellular O_2_^−^ production induced by ionomycin (0.25 μM) was determined in neutrophils without any additive (Ionomycin alone), and in neutrophils sensitized with Cmp58 alone (1 μM, 5 min) or combined with the FFA2R antagonist CATPB (100 nM). The responses were determined from the peak activities and expressed in Mcpm (mean+SD, n = 3). Statistical analyses were performed using a one-way ANOVA followed by a Dunnett’s multiple comparison test comparing the peak responses in the different scenarios described above. ns = no statistical significant difference. (**B**) Neutrophil treated (sensitized) with Cmp58 (1 μM, 5 min) were activated with ionomycin (0.25 μM) and the O_2_^−^ production was measured extracellularly (white bar) or intracellularly (black bars). The activity was determined with neutrophils incubated without or with the NADPH-oxidase inhibitor DPI (0.5 μM; 5 min). The release of O_2_^−^ was determined as the peak activities (Mcpm; (mean +SD, n=3). The statistical analysis was performed using a paired Student’s t-test comparing the peak responses induced either by absence or presence of DPI when activating with ionomycin. (**C**) The intracellular-(solid line) or extracellular-(dashed line) O_2_^−^ production was determined in neutrophils incubated (sensitized) with Cmp58 (1 μM, 5 min) and activated by ionomycin (0.25 μM). The time point for addition of ionomycin is indicated by an arrow, and one out of three independent experiments is shown **Inset:** The time to reach the peak O_2_^−^ production extracellularly (white bar) or intracellularly (black bar) was calculated using ionomycin (0.25μM) as the activating compound. The timepoints (in min) to reach peak O_2_^−^ production were determined (Mcpm; (mean + SD, n= 3).

### The FFA2Rs are selectively desensitized by ionomycin

Despite the fact that the [Ca^2+^]_i_ remained high for a long time when ionomycin was used as a tool compound to induce a rise in [Ca^2+^]_i_, the NADPH-oxidase activity was fairly rapidly terminated (see Fig 3 and 7). This suggests that the down-stream signaling becomes terminated and FFA2R is no longer sensitive to the high level of [Ca^2+^]_i_. Commonly, neutrophil responses induced by GPCR specific agonists are fairly rapidly terminated and the receptor is turned non-responsive to activating agonist recognized by the receptor, a phenomenon known as homologous receptor desensitization [13].

**Fig 7.**
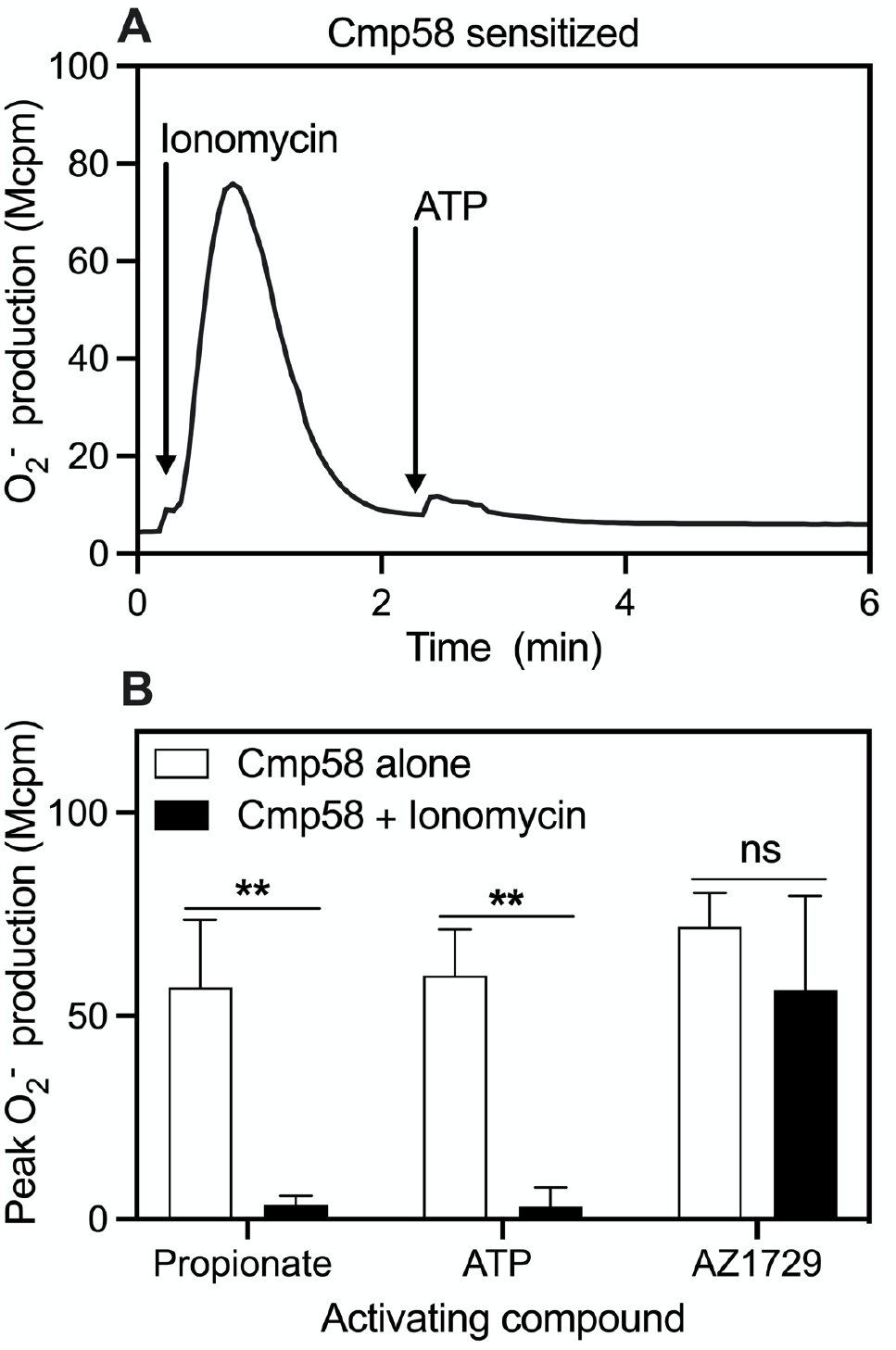
Ionomycin induced FFA2R activation of the NADPH-oxidase selective desensitizes FFA2R. (**A**) Neutrophils incubated (sensitized) with Cmp58 (1 μM, 5 min) were activated by ionomycin (0.25 μM; added at the time point indicated by the first (left) arrow). Once the response was terminated and the activity was back to baseline, the cells were activated by ATP (10 μM; added at the time point indicated by the second arrow (to the right). The production of (O_2_^−^) was followed and is expressed in Mcpm. One representative experiment out of 3 is shown. (**B**) Neutrophils incubated (sensitized) with Cmp58 (1 μM) were either activated by ionomycin (0.25 μM; black bars) or left alone. Once the response induced by ionomycin was terminated and the activity was back to baseline, these cell (black bars) as well as the non-ionomycin activated cells (white bars) were activated by propionate (25 μM), ATP (10 μM) or AZ1729 (1 μM), and the O_2_^−^ release induced by these compounds was determined. The results obtained are presented as peak O_2_^−^ production expressed as Mcpm (mean + SD, n = 3).

To determine the signaling state of FFA2R, once the ionomycin induced activation of the NADPH-oxidase has been terminated, the cells were once again activated. No activation of the NADPH-oxidase was, however, induced in these cells by the orthosteric FFA2R agonist propionate and also ATP lacked the ability to activate these cells (Fig 7A). FFA2R is, thus, turned insensitive to an orthosteric FFA2R agonist as well as to the transactivation signals generated by P2Y_2_R (Fig 7B). As mentioned, the FFA2R selective PAM AZ1729 has earlier been shown be recognized by an allosteric binding site physically/functionally separated from the binding site for Cmp58. When combined with Cmp58, this non-activating modulator was turned into a potent neutrophil activating complementary modulator (Fig 5A). The activation signals down-stream of FFAR2 when stimulated by the two interdependent complementary allosteric modulators have been shown to be biased in that, unlike for orthosteric agonists, the two modulators together triggered an activation of the NADPH-oxidase, without any concomitant rise in [Ca^2+^]_i_ [12]. The allosteric FFA2R modulator AZ1729 activated also the ionomycin “desensitized”neutrophils (Fig 7B), showing that the NADPH-oxidase activating signal can be generated by FFA2R when the receptor interacts with the two interdependent allosteric modulators Cmp58 and AZ1729 but not when either of the two agonists propionate (orthosteric FFA2R agonist) and ATP (P2Y_2_R agonist), respectively, was used to activate the cells (Fig 7B).

## Discussion

The allosteric modulator Cmp58, specific for free fatty acid receptor 2 (FFA2R), transfered ATP into a neutrophil activating compound, and this activation was FFA2R transduced and achieved without the involvement of any orthosteric FFA2R agonist. ATP, the agonist that transactivates FFA2R, is specific for the G protein coupled receptor P2Y_2_R. Obviously, the general accepted perception, that the effect of a positive allosteric receptor modulator should be restricted to an amplification of the response induced by orthosteric agonists recognized by the modulated receptor, has to be re-evaluated. This is also supported by the fact that the allosteric modulator Cmp58, turned not only ATP but also platelet activating factor (PAF) and formyl peptide receptor (FPR) agonists, into neutrophil activating compounds without the involvement of any orthosteric FFA2R agonist. These FFA2R transactivating agonists are specific for the G protein coupled PAF receptor and the FPRs 1 and 2, respectively. Referring to P2Y_2_R and the PAFR, the transactivation of FFA2R starts downstream of the Gα_q_-containing G protein coupled to the ATP/PAF receptors, and consequently the signals generated transactivate the allosterically modulated FFAR2s from the cytosolic side of the receptor expressing plasma membrane. The experimental data presented on the effects of receptor-specific antagonists, support the receptor crosstalk-signaling mechanism between P2Y_2_R and FFA2R as the molecular bases for ATP-induced activation of the neutrophil NADPH oxidase. It is clear that the Gα_q_ inhibitor YM-254890 inhibited the transactivating signals generated by ATP/P2Y_2_R and that the FFA2R transactivating signals are generated downstream of Gαq. FFA2R, thus, becomed part of the Gαq-transduced response induced by ATP when this agonist is combined with the allosteric FFA2R modulator Cmp58. Moreover, the inhibitory effect of the FFA2R antagonist CATPB on the Cmp58/ATP-induced response as well as the inability to obtain any transactivation in monocytes that lack functional FFA2Rs, were also of importance for the conclusion that FFA2R is part of the response. The transactivation signals generated downstream of the G protein has earlier only been speculated on [13], but the fact that signaling downstream of all the transactivating receptors include a rise in [Ca^2+^]_i_, opened for the possibility that this pathway was involved in the transactivation.

The transient rise in [Ca^2+^]_i_, triggered during activation of P2Y_2_R, is mediated by the Gαq-PLC-PIP_2_-IP_3_ pathway and this pathway may be activated also by β/γ subunits downstream of Gαi coupled GPCRs [21]. Irrespectively of the presice activating mechanism, the rise in [Ca^2+^]_i_ is intitiated by IP_3_ that triggers an increase in the concentration of free calcium ions cytosolicaly, ions originating from the Ca^2+^ storing endoplasmic reticulum. An emptying of the stores regulates an openening of a store operated Ca^2+^ channel in the plasma membrane allowing Ca^2+^to enter the cytosol from outside. Regarding the described receptor transactivation/crosstalk, it is important to notice that the ATP/P2Y_2_R-mediated transactivation was obtained only when the FFA2Rs were allosterically modulated. No cross-talk/transactivation was achieved with any of the receptor specific agonists in the absence of the allosteric modulator. But more importantly, the orthosteric agonist propionate could not replace the allosteric modulator in the receptor cross-talk/transactivation. The different functional outcome for binding of the allosteric modulator and the orthosteric FFA2R agonist suggests that the binding site for the modulator is outside the orthosteric binding pocket of the FFA2R. However, the effect of the FFA2R antagonist CATPB that inhibitd the activity mediated not only by propionate but also by Cmp58 suggests that the binding sites for the two FFA2R-interacting compounds (propionate and Cmp58) are very close and possibly overlap. This suggestion is supported by the results with another structurally unrelated FFA2R antagonist [20, 33] that has an inhibitory profile identical to that of CATPB.

In accordance with the proposed receptor transactivation model presented (see Fig 8), we showed that a rise in [Ca^2+^]_i_ was sufficient to activate FFA2R from inside the plasma membrane. Based on the results showing that the ATP induced increase in [Ca^2+^]_i_ was unaffected by the allosteric FFA2R modulator as well as by the FFA2R antagonist CATPB, we conclude, that the Gα_q_ coupled PLC–PIP_2_–IP_3_–Ca^2+^ signaling route downstream of P2Y_2_R is not affected by Cmp58. Instead, the data suggest that the basis for the transactivation of the Cmp58 modulated FFA2R is a changed Ca^2+^sensitivity of this receptor. It was for long generally accepted, that a receptor induced rise in [Ca^2+^]_i_ directly trigger the release of O_2_^−^ generated by the neutrophil NADPH-oxidase. This view was, however, quickly abandoned when methods were developed that made it possible to measure the concentration of ionized calcium in the cytoplasm of small cells such as neutrophil, and it was shown that the increased [Ca^2+^]_i_ induced by an ionophore such as ionomycin was in itself insufficient to trigger a release of O_2_^−^ [24]. The data presented in this study show, however, that such a release of O_2_^−^ was indirectly induced not only by ionomycin but also by thapsigargin (a drug that increases [Ca^2+^]_i_ through an inhibition of the Ca^2+^ATPase in the endoplasmic reticulum), provided that these tools to increase [Ca^2+^]_i_ were supported by allosterically modulated FFA2Rs. We have earlier shown that ionomycin induces an assembly and activation of the NADPH-oxidase localized in neutrophil granules, an activation that gives rise to a large production ROS intracellularly [34]. Since ROS have been suggested to be involved in receptor transactivation [30], the importance of the intracellular ROS production was investigated using the NADPH-oxidase inhibitor DPI, a research tool displaying a selective inhibition of the intracellular neutrophil oxidase activity [32]. The results showed that intracellular oxidase activity induced by ionomycin was not of importance for the allosteric FFA2R modulator dependent release of ROS induced by the Ca^2+^ ionophore. We also showed that, provided that FFA2R was allosterically modulated, a response was induced by several different GPCRs agonists. These results suggest that the receptor cross-talk/transactivation signals may be generated down-stream of both Gα_i_ and Gα_q_ containing G proteins and that identical intracellular cross-talk signals are generated by these receptors, signals that activate the allosterically modulated FFA2R. Regarding the role of a transient rise in [Ca^2+^]_i_ for activation of the NADPH oxidase by signals generated by GPCRs it has been shown that although ligands such as the FPR1 agonist fMLF and the PAFR agonist PAF trigger a rise in [Ca^2+^]_i_, only fMLF provokes an activation of the oxidase, suggesting that an unknown additional signal may be required [35, 36]. Despite the fact that an activation of the neutrophil NADPH-oxidase can be obtained without any rise in [Ca^2+^]_i_, the receptor transactivation data support a two-signal model in which one of the receptors involved (i.e., P2Y_2_R) is responsible for the Ca^2+^ rise and the transactivated FFA2R generates the second (unknown) signal. In this activation, the role of the signal generated by P2Y_2_R was primarily to transactivate the allosterically modulated FFA2R. In agreement with the reciprocity that is one of the characteristic parts of the allosteric GPCR modulation concept [37], the positive modulating effect of Cmp58 on the response induced by other FFA2R ligands (i.e., propionate as well as AZ1729) was basically the same when the order for addition of Cmp58 and propionate/AZ1729 was reversed and the latter were used as the sensitizing ligand. These results are, thus, in agreement with the reciprocity characterizing the effects of allosteric modulators. In contrast there was no reciprocity of the positive modulating effect of Cmp58 on the response induced by ATP, and this was true also for ionomycin. Based on the fact that the order by which the allosteric modulator and ATP/ionomycin was added to neutrophils could not be reversed, we conclude that the allosterically modulated FFA2R has to be transactivated in order for the oxidase activating signal to be generated.

**Fig 8.**
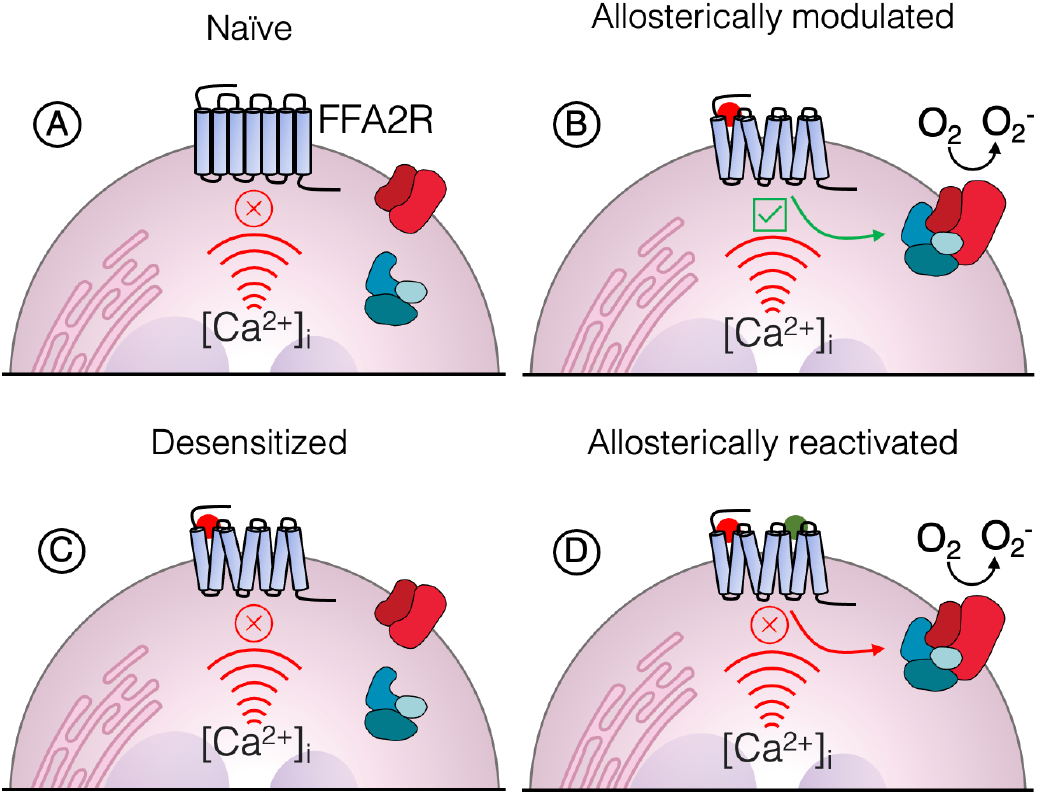
Neutrophil FFA2R activated by a rise in the cytosolic concentration of free calcium ions [Ca^2+^]_i_, and the desensitized FFA2R is allosterically reactivated. The naïve FFA2R (**A**) is not activated by an increase in [Ca^2+^]_i_ irrespectively if this increase is the result of the signaling cascade induced downstream of GPCRs such as P2Y_2_R/FPRs coupling to Gα_q_/Gα_i_. containing G proteins or by a calcium specific ionophore. In contrast, the structural change of FFA2R induced by the allosteric modulator Cmp58 (**B**) converts the receptor state that is sensitive to an increase in [Ca^2+^]_i_. The PLC–IP_3_–Ca^2+^ signaling pathway activated down-stream of the cross-talking GPCRs that couple to Gα_q_/Gα_i_. containing G proteins transactivate FFA2R from the cytosolic side of the plasma membrane. When the ionomycin induced response is terminated (**C**) the receptor is insensitive to Ca^2+^ and non-responsive (desensitized) to an addition of propionate as well as to the transactivation signal generated by ATP/P2Y_2_R, but the receptor can still be reactivated when the FFA2R modulator AZ1729 binds to it’s allosteric site on the “desensitized” receptor (**D**).

It is clear, that following termination of the ionomycin induced response, the FFA2Rs can no longer be activated by propionate or transactivated by the signal generated by ATP/P2Y_2_R. These “desinsitized” FFA2Rs can, however, still be activated and generate NADPH-oxidase activating signals. Such a response was induced when the allosteric FFA2R modulator AZ1729 was used as the co-activating ligand. The mechanisms that determine non-responsiveness/desensitization of neutrophil GPCRs such as the FFA2Rs are very complex and may be dependent on changes at the receptor level as well as at the down-stream signaling level [38–40]. But irrespective of the precise mechanism that transfers FFA2R to a biased desensitized state, the selectively desensitized receptor can still be activated. These results are in agreement with the results presented in a study using another slightly different allosteric FFA2R ligand (4-CMTB) showing that that desensitized FFA2Rs may be reactivated by a mechanism designed to safeguard FFA2R function and by that preserve the neutrophil responsiveness to othosteric FFA2R agonists [38]. The precise signals generated when AZ1729 is used to activate the cells can only be speculated on, but it is clear that an oxidase activating signal was generated also when the “biased desensitized” FFAR2s were activated by this modulating ligand. Regarding the transactivation/receptor crosstalk, several mechanisms, either at the level of separated signaling downstream of two cross-talking receptors or at the level of the participating receptors, have been described [41, 42]. Hence, it is clear that GPCRs have intracellular sites important for signaling that affect receptor functions through interactions with regulating/modulating ligands present at the cytosolic side of the plasma membrane [43]. The data presented provide, however, direct evidence for a transactivation mechanism that results in an activation of allosterically modulated FFA2Rs. This crosstalk between one receptor (i.e., P2Y_2_R) activated by its specific orthosteric agonist, and another receptor (i.e., FFA2R) that is allosterically modulated, represents a new regulatory mechanism that controls neutrophil-receptor signaling from the inside of the plasma membrane.

## Material and Methods

### Chemicals

Isoluminol, luminol, TNF-α, fMLF, ATP, PAF, diphenyleneiodonium (DPI) and propionic acid, were purchased from Sigma (Sigma Chemical Co., St. Louis, MO, USA). Dextran and Ficoll-Paque were obtained from GE-Healthcare Bio-Science (Uppsala, Sweden). Horseradish peroxidase (HRP) was obtained from Boehringer Mannheim (Mannheim, Germany). The allosteric FFA2R modulator Cmp58 ((*S*)-2-(4-chlorophenyl)-3,3-dimethyl-*N*-(5-phenylthiazol-2-yl)butanamide, the P2Y_2_R antagonist AR-C118925 {5-[[5-(2,8-dimethyl-5Hdibenzo[*α,d*]cyclohepten-5-yl)-3,4-dihydro-2-oxo-4-thioxo-1(*2H*)-pyrimidinyl]methyl]-*A*-*2H*-tetrazol-5-yl-2-furan-carboxamide},and the Ca^2+^ probes thapsigargin and ionomycin were obtained from Calbiochem-Merck Millipore (Billerica, USA). The allosteric FFAR2 modulator AZ1729 [12, 44] was from AstraZeneca (Gothenburg, Sweden). The FFA2R specific antagonist CATPB ((*S*)-3-(2-(3-chlorophenyl)acetamido)-4-(4-(trifluoromethyl)phenyl) butanoic acid used, was synthesized as described previously[19, 20]. Superoxide dismutase (SOD) and catalase were obtained from Worthington (Columbus,OH, USA), and Fura-2 AM was from Thermo Fisher Scientific (Waltham, MA, USA).

The Gα_q_ inhibitor YM-254890 was purchased from Wako Chemicals (Neuss, Germany). The FPR2 specific hexapeptide Trp-Tyr-Met-Val-Met-NH2 (WKYMVM) was synthesized and purified by HPLC by Alta Bioscience (University of Birmingham, Birmingham, United Kingdom). Subsequent dilutions of receptor ligand and other reagents were made in Krebs-Ringer Glucose phosphate buffer (KRG; 120 mM NaCl, 4.9 mM KCl, 1.7 mM KH_2_PO_4_, 8.3 mM NaH_2_PO_4_, 1.2 mM MgSO_4_, 10 mM glucose, and 1 mM CaCl_2_ in dH_2_O, pH 7.3).

### Isolation of human neutrophils/monocytes

Neutrophils were isolated from buffy coats from healthy blood donors by dextran sedimentation and Ficoll-Paque gradient centrifugation as described by Bøyum [45].

Remaining erythrocytes were removed by hypotonic lysis and the neutrophils were washed and resuspended in KRG. Human peripheral blood monocytes were isolated by the classical monocyte isolation kit (Miltenyi Biotec, #130-117-337), according to manufacturer’s instructions. To amplify the activation signals the cells were primed with TNF-α (10 ng/mL for 20 min at 37°C), and then stored on ice until use.

### Measuring NADPH-oxidase activity

An isoluminol-enhanced chemiluminescence (CL) technique was used as described [46, 47], to measure superoxide release, the precursor of production of reactive oxygen species (ROS) generated by the NADPH-oxidase. In short, the measurements were performed in a six-channel Biolumat LB 9505 (Berthold Co., Wildbad, Germany), using disposable 4-mL polypropylene tubes and a 900 μL reaction mixture containing 10^5^ neutrophils, isoluminol (0.2 μM) and HRP (4 Units/mL). The tubes were equilibrated for 5 min at 37°C, before addition of agonist (100 μL) and the light emission was recorded continuously over time. In experiments where the effects of receptor specific antagonists were determined, these were added to the reaction mixture 1–5 min before stimulation with control neutrophils incubated under the same condition but in the absence of antagonist run in parallel for comparison.

A luminol-enhanced chemiluminescence (CL) technique was used to measure intracellular production of ROS generated by the NADPH-oxidase activity in principle as described above but, in the disposable 4-mL polypropylene tubes with a 900 μL reaction mixture, the membrane permeable amplifier luminol (0.2 μM) together with superoxide dismutase (SOD; 200Units) and catalase (2000 Units) replaced isoluminol and HRP (for details see [46, 47]).

### Calcium mobilization

Neutrophils (2 × 10^7^/ml) were loaded with Fura-2 AM (2 μM) in KRG containing 0.1% BSA for 30 minutes in darkness at room temperature before washing and suspension in KRG at a density of 2×10^7^/ml. Calcium measurements were carried out in a Perkin Elmer fluorescence spectrophotometer (LC50), with excitation wavelengths of 340 nm and 380 nm, an emission wavelength of 509 nm, and slit widths of 5 nm and 10 nm, respectively. The transient rise in intracellular calcium is presented as the ratio of fluorescence intensities (340 nm / 380 nm) detected.

### Statistical analysis

Statistical calculations were performed in GraphPad Prism 9.3 (Graphpad Software, San Diego, CA, USA). Which specific statistical tests that are performed are stated in the relevant figure legend. A *p*-value < 0.05 was regarded as statistically significant difference and is indicated by \**p* < 0.05, \*\**p* < 0.01 \*\*\**p* < 0.001. Even though the results in the figures are presented as percent of control values, the statistics have been calculated using the raw data.

### Ethics Statement

In this study, conducted at the Sahlgrenska Academy in Sweden, buffy coats obtained from the blood bank at Sahlgrenska University Hospital, Gothenburg, Sweden have been used. According to the Swedish legislation section code 4§ 3p SFS 2003:460 (Lag om etikprövning av forskning som avser människor), no ethical approval was needed since the buffy coats were provided anonymously and could not be traced back to a specific donor.

## Abbreviations

CATPB: (*S*)-3-(2-(3-chlorophenyl)acetamido)-4-(4-(trifluoromethyl)phenyl)butanoic acid an FFA2R antagonist
[Ca^2+^]_i_: intracellular concentration of free calcium ions
CL: chemiluminescence
Cmp58: an allosteric FFA2R modulator
FFA2R: free fatty acid receptor 2
DAMP: danger associated molecular pattern
DPI: diphenyleneiodonium
FPR: formyl peptide receptor
GPCR: G protein coupled receptor
HRP: horse radish peroxidase
ns: no statistical significant difference
KRG: Krebs-Ringer Glucose phosphate buffer
O_2_^−^: superoxide
PAFR: receptor for platelet activating factor
PAM: positive allosteric modulator
PAMP: pathogen associated molecular pattern
P2Y_2_R: receptor for ATP
ROS: reduced/reactive oxygen species
TNF-α: tumor necrosis factor α

## Acknowledgements

The authors thank the members of the Phagocyte Research Group at the Sahlgrenska Academy, University of Gothenburg for critically discussing the results and the manuscript. The experimental work performed by Linda Bergqvist (University of Gothenburg) is also gratefully acknowledged. The FFA2R antagonist CATPB was a generous gift from Trond Ulven (Odense University, Denmark), and the allosteric FFA2R modulator AZ1729 was kindly provided by Kenneth Granberg (BioPharmaceuticals R&D, AstraZeneca, Gothenburg, Sweden).

## Funding statement

The work was supported by the Swedish Medical Research Council (HF, 2018-02848) and the Swedish state under the ALF-agreement (CD, ALFGBG 72510; HF, ALFGBG78150), the Clas Groschinskys Memorial Fund (MS M21146), the Åke Wiberg Foundation (MS M21-0025), the Sahlgrenska International Starting Grant (MS), the Magnus Bergwall Foundation (MS 2021-04110). The sponsors did not have any role in any part of the study.

## Author contribution

S. L. and Y.W. performed research and analyzed data.

H.F., M.S., C.D. supervised the research.

C. D. planned the work together with S.L. and also analyzed data and wrote the manuscript that was edited by all authors.

## Conflict of interest

The authors declare that they have no conflict of interest with the content of this article.

## Notes

### Competing Interest Statement

The authors have declared no competing interest.

## References

[1] P.M. Mortimer, S.A. Mc Intyre, D.C. Thomas, Beyond the Extra Respiration of Phagocytosis: NADPH Oxidase 2 in Adaptive Immunity and Inflammation, Front Immunol 12 (2021) 733918.

[2] T. Lammermann, W. Kastenmuller, Concepts of GPCR-controlled navigation in the immune system, Immunol Rev 289(1) (2019) 205–231.

[3] M. Metzemaekers, M. Gouwy, P. Proost, Neutrophil chemoattractant receptors in health and disease: double-edged swords, Cell Mol Immunol 17(5) (2020) 433–450.

[4] D.A. Sykes, L.A. Stoddart, L.E. Kilpatrick, S.J. Hill, Binding kinetics of ligands acting at GPCRs, Mol Cell Endocrinol 485 (2019) 9–19.

[5] J.S. Smith, R.J. Lefkowitz, S. Rajagopal, Biased signalling: from simple switches to allosteric microprocessors, Nat Rev Drug Discov 17(4) (2018) 243–260.

[6] P. Kolb, T. Kenakin, S.P.H. Alexander, M. Bermudez, L.M. Bohn, C.S. Breinholt, M. Bouvier, S.J. Hill, E. Kostenis, K.A. Martemyanov, R.R. Neubig, H.O. Onaran, S. Rajagopal, B.L. Roth, J. Selent, A.K. Shukla, M.E. Sommer, D.E. Gloriam, Community guidelines for GPCR ligand bias: IUPHAR review 32, Br J Pharmacol (2022).

[7] A.K. Shukla, Emerging paradigms in activation, signaling, and regulation of G protein-coupled receptors, FEBS J 288(8) (2021) 2458–2460.

[8] L.M. Slosky, M.G. Caron, L.S. Barak, Biased Allosteric Modulators: New Frontiers in GPCR Drug Discovery, Trends Pharmacol Sci 42(4) (2021) 283–299.

[9] M. Grundmann, E. Bender, J. Schamberger, F. Eitner, Pharmacology of Free Fatty Acid Receptors and Their Allosteric Modulators, Int J Mol Sci 22(4) (2021).

[10] G. Milligan, B. Shimpukade, T. Ulven, B.D. Hudson, Complex Pharmacology of Free Fatty Acid Receptors, Chem Rev 117(1) (2017) 67–110.

[11] S. Lind, A. Holdfeldt, J. Martensson, K.L. Granberg, H. Forsman, C. Dahlgren, Multiple ligand recognition sites in free fatty acid receptor 2 (FFA2R) direct distinct neutrophil activation patterns, Biochem Pharmacol 193 (2021) 114762.

[12] S. Lind, A. Holdfeldt, J. Martensson, M. Sundqvist, T.P. Kenakin, L. Bjorkman, H. Forsman, C. Dahlgren, Interdependent allosteric free fatty acid receptor 2 modulators synergistically induce functional selective activation and desensitization in neutrophils, Biochim Biophys Acta Mol Cell Res 1867(6) (2020) 118689.

[13] C. Dahlgren, A. Holdfeldt, S. Lind, J. Martensson, M. Gabl, L. Bjorkman, M. Sundqvist, H. Forsman, Neutrophil Signaling That Challenges Dogmata of G Protein-Coupled Receptor Regulated Functions, ACS Pharmacol Transl Sci 3(2) (2020) 203–220.

[14] C. Dahlgren,Lind, S.,Mårtensson, J.,Björkman, L.,Wu, Y.,Sundqvist, M and H.Forsman., G PROTEIN COUPLED PATTERN RECOGNITION RECEPTORS EXPRESSED IN NEUTROPHILS: Recognition, activation/modulation, signaling and receptor regulated functions Immunological reviews In press (2022).

[15] S. Lind, A. Holdfeldt, J. Martensson, M. Sundqvist, L. Bjorkman, H. Forsman, C. Dahlgren, Functional selective ATP receptor signaling controlled by the free fatty acid receptor 2 through a novel allosteric modulation mechanism, FASEB J 33(6) (2019) 6887–6903.

[16] L.E. Kilpatrick, S.J. Hill, Transactivation of G protein-coupled receptors (GPCRs) and receptor tyrosine kinases (RTKs): Recent insights using luminescence and fluorescence technologies, Curr Opin Endocr Metab Res 16 (2021) 102–112.

[17] S. Palanisamy, C. Xue, S. Ishiyama, S.V. Naga Prasad, K. Gabrielson, GPCR-ErbB transactivation pathways and clinical implications, Cell Signal 86 (2021) 110092.

[18] W. Wang, Y. Qiao, Z. Li, New Insights into Modes of GPCR Activation, Trends Pharmacol Sci 39(4) (2018) 367–386.

[19] B.D. Hudson, M.E. Due-Hansen, E. Christiansen, A.M. Hansen, A.E. Mackenzie, H. Murdoch, S.K. Pandey, R.J. Ward, R. Marquez, I.G. Tikhonova, T. Ulven, G. Milligan, Defining the molecular basis for the first potent and selective orthosteric agonists of the FFA2 free fatty acid receptor, J Biol Chem 288(24) (2013) 17296–312.

[20] E. Sergeev, A.H. Hansen, S.K. Pandey, A.E. MacKenzie, B.D. Hudson, T. Ulven, G. Milligan, Non-equivalence of Key Positively Charged Residues of the Free Fatty Acid 2 Receptor in the Recognition and Function of Agonist Versus Antagonist Ligands, J Biol Chem 291(1) (2016) 303–17.

[21] T. Kawakami, W. Xiao, Phospholipase C-beta in immune cells, Adv Biol Regul 53(3) (2013) 249–57.

[22] N. Demaurex, S. Saul, The role of STIM proteins in neutrophil functions, J Physiol 596(14) (2018) 2699–2708.

[23] C. Dahlgren, Difference in extracellular radical release after chemotactic factor and calcium ionophore activation of the oxygen radical-generating system in human neutrophils, Biochim Biophys Acta 930(1) (1987) 33–8.

[24] T. Pozzan, D.P. Lew, C.B. Wollheim, R.Y. Tsien, Is cytosolic ionized calcium regulating neutrophil activation?, Science 221(4618) (1983) 1413–5.

[25] D. Ribeiro, M. Freitas, S. Rocha, J. Lima, F. Carvalho, E. Fernandes, Calcium Pathways in Human Neutrophils-The Extended Effects of Thapsigargin and ML-9, Cells 7(11) (2018).

[26] S. Lind, D.O. Hoffmann, H. Forsman, C. Dahlgren, Allosteric receptor modulation uncovers an FFA2R antagonist as a positive orthosteric modulator/agonist in disguise, Cell Signal 90 (2022) 110208.

[27] L. Bjorkman, J. Martensson, M. Winther, M. Gabl, A. Holdfeldt, M. Uhrbom, J. Bylund, A. Hojgaard Hansen, S.K. Pandey, T. Ulven, H. Forsman, C. Dahlgren, The Neutrophil Response Induced by an Agonist for Free Fatty Acid Receptor 2 (GPR43) Is Primed by Tumor Necrosis Factor Alpha and by Receptor Uncoupling from the Cytoskeleton but Attenuated by Tissue Recruitment, Mol Cell Biol 36(20) (2016) 2583–95.

[28] M. Sundqvist, K. Christenson, A. Holdfeldt, M. Gabl, J. Martensson, L. Bjorkman, R. Dieckmann, C. Dahlgren, H. Forsman, Similarities and differences between the responses induced in human phagocytes through activation of the medium chain fatty acid receptor GPR84 and the short chain fatty acid receptor FFA2R, Biochim Biophys Acta Mol Cell Res 1865(5) (2018) 695–708.

[29] C. Dahlgren, A. Karlsson, Ionomycin-induced neutrophil NADPH oxidase activity is selectively inhibited by the serine protease inhibitor diisopropyl fluorophosphate, Antioxid Redox Signal 4(1) (2002) 17–25.

[30] R. Mohamed, R. Janke, W. Guo, Y. Cao, Y. Zhou, W. Zheng, H. Babaahmadi-Rezaei, S. Xu, D. Kamato, P.J. Little, GPCR transactivation signalling in vascular smooth muscle cells: role of NADPH oxidases and reactive oxygen species, Vasc Biol 1(1) (2019) R1–R11.

[31] B.V. O’Donnell, D.G. Tew, O.T. Jones, P.J. England, Studies on the inhibitory mechanism of iodonium compounds with special reference to neutrophil NADPH oxidase, Biochem J 290 (Pt 1)(1993) 41–9.

[32] A. Buck, F.P. Sanchez Klose, V. Venkatakrishnan, A. Khamzeh, C. Dahlgren, K. Christenson, J. Bylund, DPI Selectively Inhibits Intracellular NADPH Oxidase Activity in Human Neutrophils, Immunohorizons 3(10) (2019) 488–497.

[33] F. Namour, R. Galien, T. Van Kaem, A. Van der Aa, F. Vanhoutte, J. Beetens, G. Van’t Klooster, Safety, pharmacokinetics and pharmacodynamics of GLPG0974, a potent and selective FFA2 antagonist, in healthy male subjects, Br J Clin Pharmacol 82(1) (2016) 139–48.

[34] C. Dahlgren, A. Johansson, H. Lundqvist, O.W. Bjerrum, N. Borregaard, Activation of the oxygen-radical-generating system in granules of intact human neutrophils by a calcium ionophore (ionomycin), Biochim Biophys Acta 1137(2) (1992) 182–8.

[35] S. Brechard, E.J. Tschirhart, Regulation of superoxide production in neutrophils: role of calcium influx, J Leukoc Biol 84(5) (2008) 1223–37.

[36] R. Foyouzi-Youssefi, F. Petersson, D.P. Lew, K.H. Krause, O. Nusse, Chemoattractant-induced respiratory burst: increases in cytosolic Ca2+ concentrations are essential and synergize with a kinetically distinct second signal, Biochem J 322 (Pt 3) (1997) 709–18.

[37] A. Christopoulos, T. Kenakin, G protein-coupled receptor allosterism and complexing, Pharmacol Rev 54(2) (2002) 323–74.

[38] R. Frei, J. Nordlohne, U. Huser, S. Hild, J. Schmidt, F. Eitner, M. Grundmann, Allosteric targeting of the FFA2 receptor (GPR43) restores responsiveness of desensitized human neutrophils, J Leukoc Biol 109(4) (2021) 741–751.

[39] S. Rajagopal, S.K. Shenoy, GPCR desensitization: Acute and prolonged phases, Cell Signal 41 (2018) 9–16.

[40] N. Sun, K.M. Kim, Mechanistic diversity involved in the desensitization of G protein-coupled receptors, Arch Pharm Res 44(4) (2021) 342–353.

[41] L. Prezeau, M.L. Rives, L. Comps-Agrar, D. Maurel, J. Kniazeff, J.P. Pin, Functional crosstalk between GPCRs: with or without oligomerization, Curr Opin Pharmacol 10(1) (2010) 6–13.

[42] W. Stallaert, E.T. van der Westhuizen, A.M. Schonegge, B. Plouffe, M. Hogue, V. Lukashova, A. Inoue, S. Ishida, J. Aoki, C. Le Gouill, M. Bouvier, Purinergic Receptor Transactivation by the beta2-Adrenergic Receptor Increases Intracellular Ca(2+) in Nonexcitable Cells, Mol Pharmacol 91(5) (2017) 533–544.

[43] N.V. Ortiz Zacarias, E.B. Lenselink, I.J. Ap, T.M. Handel, L.H. Heitman, Intracellular Receptor Modulation: Novel Approach to Target GPCRs, Trends Pharmacol Sci 39(6) (2018) 547–559.

[44] D. Bolognini, C.E. Moss, K. Nilsson, A.U. Petersson, I. Donnelly, E. Sergeev, G.M. Konig, E. Kostenis, M. Kurowska-Stolarska, A. Miller, N. Dekker, A.B. Tobin, G. Milligan, A Novel Allosteric Activator of Free Fatty Acid 2 Receptor Displays Unique Gi-functional Bias, J Biol Chem 291(36) (2016) 18915–31.

[45] A. Boyum, D. Lovhaug, L. Tresland, E.M. Nordlie, Separation of leucocytes: improved cell purity by fine adjustments of gradient medium density and osmolality, Scand J Immunol 34(6) (1991) 697–712.

[46] C. Dahlgren, H. Bjornsdottir, M. Sundqvist, K. Christenson, J. Bylund, Measurement of Respiratory Burst Products, Released or Retained, During Activation of Professional Phagocytes, Methods Mol Biol 2087 (2020) 301–324.

[47] H. Lundqvist, C. Dahlgren, Isoluminol-enhanced chemiluminescence: a sensitive method to study the release of superoxide anion from human neutrophils, Free Radic Biol Med 20(6) (1996) 785–92.

